# Biopolymers can be used for consolidation of lunar and Martian regoliths

**DOI:** 10.1101/2025.01.03.631200

**Authors:** Nitin Gupta, Aloke Kumar, Koushik Viswanathan

## Abstract

Creating long-term extraterrestrial habitats to support continuous human occupation requires structures that can bear load, while remaining resilient to large temperature fluctuations. Exploiting locally available regolith is part of a paradigm termed *in situ* resource utilization (ISRU) that advocates minimal dependence on terrestrial supplies via local resource exploitation for construction. In this work, we report on a consolidation method for making load-bearing structures out of lunar and Martian regolith simulants using biopolymer binders. We quantify the compressive and tensile strength of the final brick-like structures, termed biopolymer regolith composites, and investigate their dependence on biopolymer concentration and chemistry. A pressure-assisted die casting process is developed that can produce bricks with compressive strengths of ∼ 15 MPa and tensile strengths of around 2 MPa, from regolith simulants. We finally report on the performance of these bricks under large temperature and pressure variations and find that their strengths remain unchanged. We believe that this makes biopolymer regolith composite bricks suitable candidates for use in harsh extraterrestrial environments, while adhering to the guidelines of the ISRU paradigm.

**Keyword:** Lunar regolith; Martian regolith; Space habitation; Extraterrestrial construction; Biopolymers; Regolith simulant

## 1 Introduction

A key challenge in creating long-term extraterrestrial habitats to support continuous human occupation lies in building structures that can bear load [1–3], while simultaneously offering protection from radiation and large temperature fluctuations [4, 5]. The most abundant material for the construction of these habitats on lunar and Mars surfaces is locally available regolith. Exploiting regolith is one part of a larger paradigm termed *in situ* resource utilization (ISRU) [6–10]. ISRU seeks to minimize dependence on terrestrial supplies and advocates the use of local resources, such as solar energy, regolith, and water, where applicable, as construction constituents [11, 12].

Consolidating regolith into fundamental building blocks—akin to terrestrial bricks—requires the use of a suitable binding process and/or agent. It is now well known that both Martian and lunar regolith exhibit several similarities with terrestrial soil [13,14]. Specifically, lunar regolith can be likened to sandy soil due to its near complete lack of inherent binding mineral (e.g., clay) as well as its poor water retention capability. On the other hand, Martian regolith resembles clayey soil more closely, possessing inherent binding materials such as hydrated silicas or gypsum [15, 16]. To mimic the behaviour of actual regolith, of which we have very few physical samples, several research groups around the world have developed suitable regolith simulants to reproduce compositions from different lunar or Martian sites. Some examples include Johnson Space Center (JSC) lunar regolith simulant [17], Lunar soil simulant by ISRO Sittampundi Anorthosite Complex (LSS-ISAC) [18], Lunar highland simulant (LHS), Lunar mare dust simulant (LMS), Mars global simulant (MGS) from Exolith Labs [16, 19], Korea Lunar Simulant (KLS) [20], and Fuji Japan Simulant (FJS) [21]. The possibility of using water for consolidating many of these regolith simulants has been explored both in the context of microbe-induced consolidation [22–24] as well as conventional sintering [25, 26]. It is also worth mentioning that more recently, waterless consolidation methods have begun to emerge as competitive processes for making bricks in certain simulants [27, 28].

Further pursuing the analogy with terrestrial soil, it is now known that an important class of natural binder materials, such as clay, bamboo, and related plant-based polysaccharides [29–31] can be used to bind terrestrial soil into load bearing consolidates. A large fraction of these natural binders are biopolymers that are also useful for sustaining plant life. Using such binders, *in lieu* of more conventional consolidation processes, such as kiln sintering, is advantageous because it is simpler and much more energy efficient, making it perfectly aligned with the ISRU paradigm. In fact, recent work on terrestrial soils has proposed the use of biopolymers, such as guar and xanthan gums, as binders, resulting in compressive strengths of up to 8 MPa [32–34]. Similarly, with extraterrestrial regolith simulants, biopolymers like Bovine Serum Albumin (BSA) [35–37], and human serum albumin (HSA) [24], etc., was used to create biopolymer soil bound composites (BSC) [38]. For the construction of space habitats, the strength requirements are quite low compared with terrestrial brick. The bricks manufactured using conventional kiln-based construction typically exhibit compressive strength of 20-25 MPa; however, due to reduced gravity, actual strength requirements on lunar and Martian surfaces are actually only 3-4 MPa and 6-7 MPa, respectively.

In the present study, we examine the consolidation potential of regolith simulants using minimal quantities of biopolymers, leading to consolidates that we term Biopolymer Regolith Composites (BRCs). We establish a process for producing BRC bricks with consistent strength and analyze how they fail under sustained loading. Failure is quantified using digital image correlation (DIC) techniques and by tracking the development and evolution of cracks within the material. Additionally, we investigate the impact of varying temperature conditions on BRCs and evaluate the degradation of biopolymers under these conditions. Our objective is to establish a scalable methodology that largely utilizes *in situ* materials for construction, with a target compressive strength of 15-20 MPa, comparable to that of conventional kiln-fired bricks on Earth.

The manuscript is organized as follows: materials used, corresponding process protocol for consolidation of BRCs, and various characterization techniques are described in Sec. 2. Measurement of BRC strength and its dependence on various process parameters is quantified in Sec. 3. Finally, we assess the sustainability of the produced bricks under simulated thermal and pressure variations. A discussion of our results with some concluding remarks is presented in Sec. 4.

## 2 Materials and Methods

### 2.1 Regolith simulants and biopolymers

We used regolith simulants procured from Space Research Technologies (Exolith labs), USA [16,19] for our experiments. Consolidation studies were performed on four different types of simulants: 2 lunar (Lunar highland simulant — LHS-1 and Lunar maredust simulant — LMS-1), and 2 Martian (Mars Global simulant — MGS-1 and Mars Global simulant sulfate — MGS-1S). The compositional details and mean particle size (⟨d*_p_*⟩) of these simulants are provided in Tables 1 and 2.

**Table 1:** Mineral composition of lunar regolith simulants.

| Minerals | LHS-1 | LMS-1 |
| --- | --- | --- |
| Anorthosite | 74.4% | 19.8% |
| Glass-rich basalt | 24.7% | 32.0% |
| Olivine | 0.2% | 11.1% |
| Pyroxene | 0.3% | 32.8% |
| Ilmenite | 0.4% | 4.3% |
| $\langle d_p \rangle$ ($\mu\text{m}$) | 88 | 63 |

**Table 2:** Mineral composition of Martian regolith simulants.

| Minerals | MGS-1 | MGS-1S |
| --- | --- | --- |
| Gypsum | - | 40% |
| Plagioclase | 27.1% | 16.4% |
| Glass-rich basalt | 22.9% | 13.7% |
| Pyroxene | 20.3% | 12.2% |
| Olivine | 13.7% | 8.2% |
| Mg-sulfate | 4.0% | 2.4% |
| Ferrihydrite | 3.5% | 2.1% |
| Hydrated silica | 3.0% | 1.8% |
| $\langle d_p \rangle$ ($\mu\text{m}$) | 90 | 119 |

Three different polysaccharide biopolymers were used for binding the regolith in this study, *viz.* guar gum (GG), xanthan gum (XG), and acacia gum (AG), procured from Urban Platter, India. GG is a galactomannan polysaccharide characterized by a mannose backbone with galactose side groups, with high molecular weight ranging from 2,00,000 to 10,00,000 [33, 39]. XG is a heteropolysaccharide composed of repeating units of glucose, mannose, and glucuronic acid in a molar ratio of approximately 2:2:1, with typical molecular weight around 6,00,000; some fractions may exceed 10,00,000 [40]. AG primarily consists of arabinogalactan, a complex polysaccharide made up of D-galactose units with side chains of D-glucuronic acid and L-arabinose or L-rhamnose, exhibiting a molecular weight ranging from 2,00,000 to 6,00,000. GG and XG are typically produced as particles of size ∼ 180 µm, with some finer grades reaching ∼ 75 µm. In contrast, AG particles typically have larger sizes ∼ 200 − 500 µm.

### 2.2 Simple casting (SC) and pressure-assisted die casting (PADC) for producing biopolymer regolith composites

Consolidation of regolith simulant using biopolymers to produce biopolymer regolith composite (BRCs) was investigated using two independent protocols, termed simple casting (SC) and pressure-assisted die casting (PADC), see Figure 1. The starting point for both protocols was the same—a mixture of regolith simulant (100g), polysaccharide gum powder of varying quantity (0 to 2% w/w of regolith simulant), and DI water (15ml) was first prepared. The regolith was thoroughly mixed with the gum in powder form, after which water was added, and the resulting mixture was stirred well.

**Figure 1:**
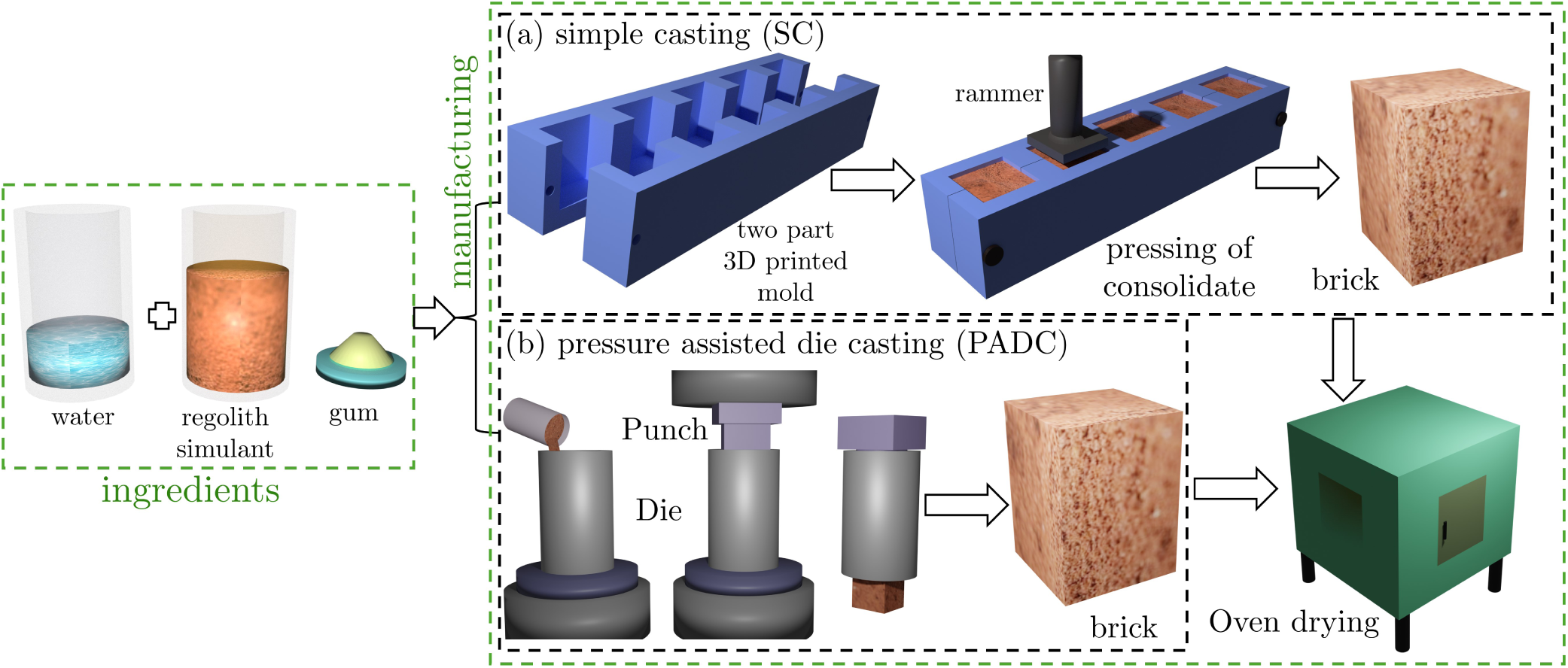
Schematic showing the method for preparing biopolymer regolith composite (BRC) bricks. Row (a) shows the simple casting (SC) method wherein a two-part mold is used, and the pre-mixed material is filled and hand-rammed. Row (b) shows the pressure-assisted die casting (PADC) method for producing BRC bricks. At the end of either process, the mold/die is oven-dried at 60*^◦^* for 12 hours.

In the simple casting (SC) method, the mixture was filled into a two-part plastic mold, whose inner surfaces were covered with a transparent plastic sheet to ease sample removal. Final BRC consolidates produced from this mold had dimensions 20×20×20 mm^3^, as shown in figure 1(a). Specimens were hand rammed and taken to oven dry at 60*^◦^*C for 12 hours. Specimens were finally polished using sandpaper to obtain a flat surface for subsequent mechanical testing.

In the pressure-assisted die casting (PADC) method, the pre-fill mixture was first compacted using a hydraulic press under a load of 5 tons and die-cast into the desired shape (cubical block 15×15×15 mm^3^, or cylindrical disc of diameter 12 mm with thickness 6-7mm), as shown in figure 1(b). The specimens were then dried in an oven at a temperature of 60*^◦^*C for 12 hours. This post-removal drying was necessary to ensure the effective removal of moisture from the consolidated.

### 2.3 Mechanical testing

The strength of consolidated BRCs was quantified by measuring their unconfined compressive strength (UCS, ASTM C109) and tensile strength (Brazilian disc configuration ASTM D3967). Both these tests were performed on a universal testing machine (UTM, Instron 5697), with 30kN (compression) and 5kN (tension) load cells under quasi-static conditions (loading rate 0.5 mm/min). The specimen dimensions were smaller than ASTM standards because of practical constraints but satisfied the plane stress condition for compression and tensile testing.

The compressive strength was calculated as 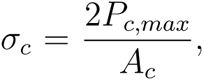 where *P_c,max_* is the maximum load at failure, and A*_c_* is the initial cross-sectional area of the specimen. The tensile strength was calculated as 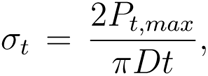 where *P_t,max_* is the maximum load at failure, and D and t are the initial diameter and the thickness of the specimen, respectively [41]. Both measurements were averaged over four specimens, with standard deviation reported in the corresponding error bars.

To quantify failure under loading, crack paths were identified and corresponding strains were evaluated using digital image correlation (DIC) analysis. For this purpose, digital images were recorded using a DSLR camera (Nikon D850) at 25 FPS and analyzed frame-by-frame using the Ncorr software package [42] with MATLAB version 2021a. The need to ensure planar deformation and through-thickness cracks (required for DIC) meant that these analyses could only be performed for the tensile samples. The cubical blocks used for compression tests inevitably failed because of multiple cracks developing through the thickness and out of the viewing plane of the camera.

## 3 Results

We present the results of various process and consolidation parameters on the final mechanical properties of biopolymer regolith composites (BRCs). The role of different biopolymer binders in determining BRC strength is also quantified, and failure mechanisms under load application are evaluated.

### 3.1 Strength of simple (SC) and pressure-assisted die cast (PADC) bricks

Compressive strengths (σ*_c_*) of BRC bricks produced via simple casting (SC) and pressure-assisted die casting (PADC) of lunar simulants LHS-1, LMS-1, and Martian simulants MGS-1 and MGS-1S, are reproduced in Fig. 2. In the left half (red) of the plot, samples were cast using only water and no biopolymer binder, while the right half (purple) are samples with 1% w/w of guar gum (GG). The same markers are used for LHS-1 (□), LMS-1 (◦), MGS-1 (△) and MGS-1S (⋆) in both halves.

**Figure 2:**
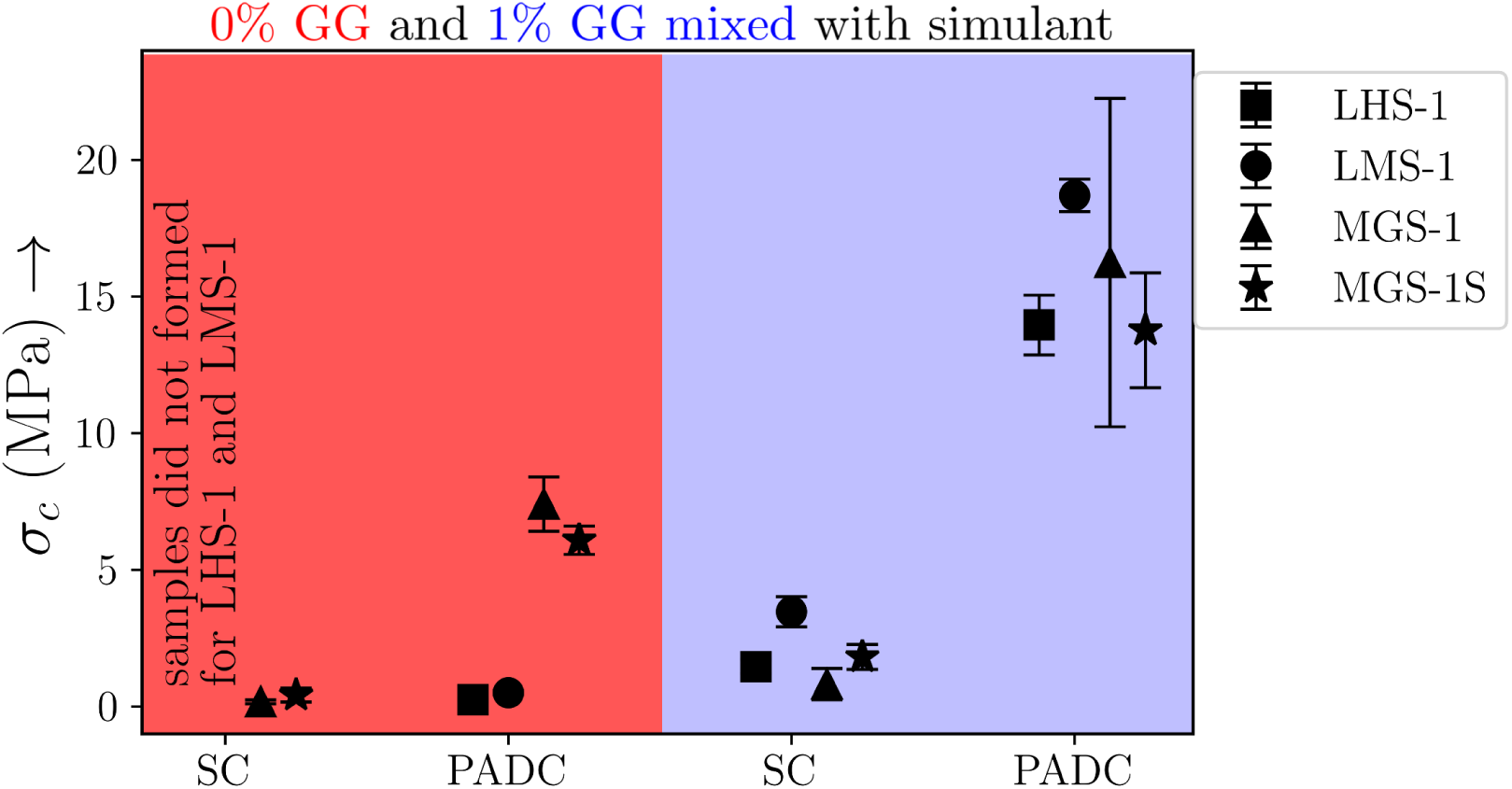
Compressive strengths σ*_c_* of regolith simulants with the SC/PADC protocols as well as with (red) and without (blue) biopolymer binders (1% w/w guar gum). The average strength was obtained from four sample measurements, with error bars representing one standard deviation.

When no binders are used, it is clear that samples produced via SC were incapable of bearing any load. In fact, it was not even possible to obtain consolidated samples with LHS-1 and LMS-1. On the other hand, weakly consolidated samples were obtained from MGS-1 and MGS-1S even in the absence of biopolymer binders, owing to the presence of inherent, but weak, binders in the simulant itself (1.8-3% hydrated silica). With the PADC process, a significant increase was observed in the compressive strengths of all four regolith simulants, with the largest seen for the Martian simulant samples. The hydrated silica (C-H-S) in these simulants effectively binds loose soil particles under high pressure and moisture conditions [43,44]. However, despite this increase in compressive strengths of over 5 MPa, the resulting samples were water soluble and would degrade within minutes when put in water.

When combined with 1% w/w of guar gum, both the compressive strength and final water resistance were significantly enhanced. All four simulants could be consolidated with the SC protocol but showed compressive strengths of not more than a few MPa. On the other hand, the PADC process produced bricks with σ*_c_* ∼ 15 MPa. This is to be expected since it is known, for instance, that biopolymers, especially polysaccharides, can form stable aggregates with terrestrial soils [45, 46]. However, the mechanisms and effectiveness of the binding are different when comparing soils with and without inherent binders [30, 32–34]. In the former case (as with MGS-1 and MGS-1S), polysaccharides predominantly adhere via weak hydrogen bonds between the carboxyl group and the hydroxyl group of the polysaccharide molecules and cations in the soil. This interaction facilitates the formation of bridges between binding mineral particles, forming microaggregates (size < 250 microns) and macroaggregates (size > 250 microns). Conversely, when no inherent binders are present (as with LHS-1 and LMS-1), polysaccharides create a cross-linked three-dimensional network, resulting in inter-particle cohesion.

Having established the importance of the polymer binder (GG), we now investigate the large increase in strength of bricks produced via PADC vis-á-vis the SC process. This can be attributed to the reduction in part porosity under the application of pressure. Optical micrograph images of samples produced via SC and PADC (1% GG) are reproduced in Fig. 3 for LHS-1 simulant. While the overall porosity (dark) is evidently lesser in the PADC produced part (panel b), the inset to these two panels shows the difference more dramatically. It is well-known that the compressive strength of concrete-like materials primarily depends on the porosity of the structure. In this case, let us consider a scenario where the overall porosity of bricks is nominally 25% [25] under PADC (measured using mercury-intrusion porosimetry) after compaction under a hydraulic press. Using the exponential relationship between porosity and strength given in Ref. [26], we estimate that the porosity in the SC case is nearly 50%.

**Figure 3:**
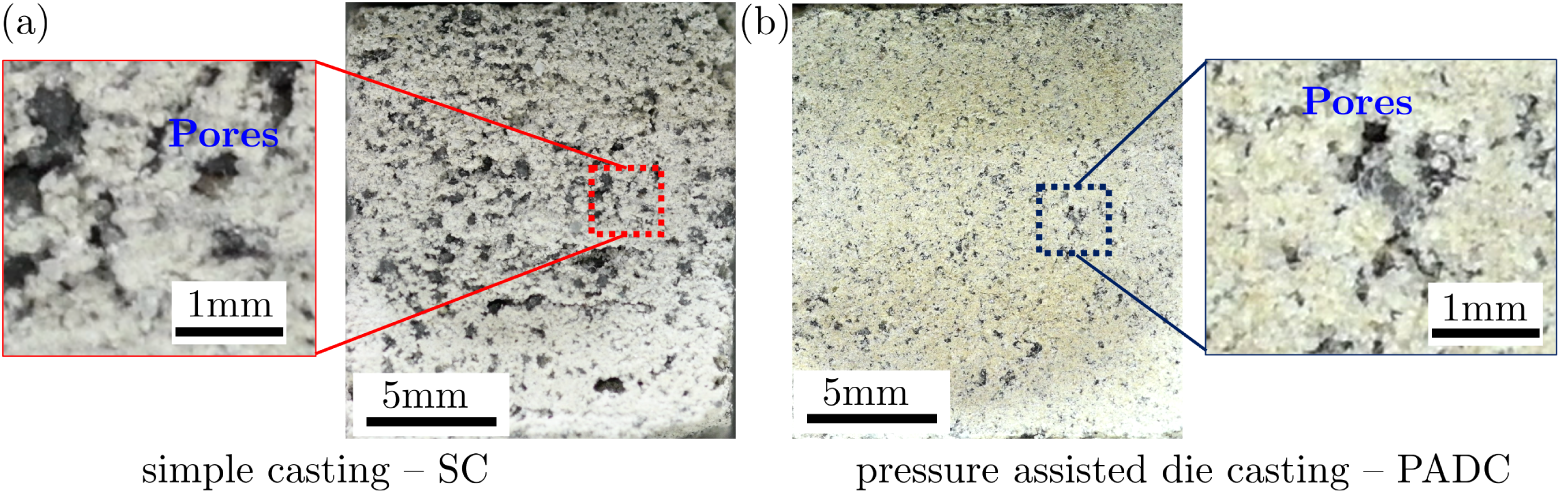
Optical image showing LHS-1 based consolidated BRC brick, with 1% w/w guar gum. (a) The sample was obtained via simple casting (SC) and (b) pressure-assisted die casting (PADC). The inset to each image clearly shows the lowered porosity in the PADC case due to the applied compaction pressure.

### 3.2 Does the quantity of gum matter?

The second question that we attempt to answer pertains to the optimal concentration of biopolymer in the final BRC. For this, guar gum (GG) content varied systematically, and corresponding compressive and tensile tests were performed. Since both lunar simulants (LMS-1 and LHS-1) and Martian simulants (MGS-1S, MGS-1) demonstrated similar behavior among themselves, we only report results for LHS-1 and MGS-1 henceforth, unless specified otherwise. Further, only results with the PADC process are reported since the strengths obtained are superior to those from the SC process.

Typical load-displacement graphs for compression (P*_c_*, red) and tension (P*_t_*, blue) are shown in Fig. 4(a). The compressive (tensile) strength σ*_c_*(σ*_t_*) was obtained from the corresponding peak P*_c,max_* (P*_t,max_*). Insets to this figure show the occurrence of cracks at the maximum load; it is clear from them that the tensile specimens (blue) failed via a single through-thickness crack (yellow arrows), while compressive specimens (red) failed via multiple crack propagation through the sample thickness (yellow arrows).

**Figure 4:**
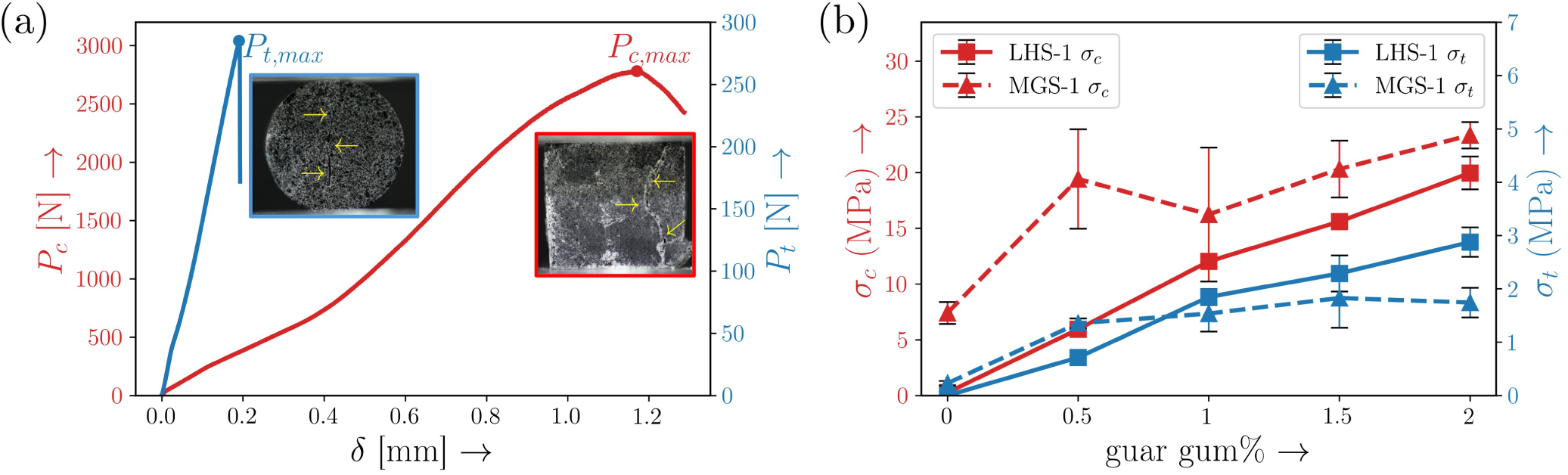
Compressive and tensile strengths of BRC bricks and their variation with binder concentration. (a) Typical load-displacement curves are used to determine the compressive and tensile strengths of BRC bricks. The inset shows a crack pattern at the point of failure, corresponding to the maximum load in tension or compression. (b) Variation in compressive (σ*_c_*) and tensile (σ*_t_*) strengths as a function of guar gum concentration. Data is for specimens (LHS-1 or MGS-1) made using the PADC process (5 tons load).

Variation in σ*_c_* (red) and σ*_t_* (blue) with GG concentration is shown in Fig. 4(b). It is clear at once that increased GG concentration leads to higher tensile and compressive strengths for both LHS-1 and MGS-1. Initially, a nearly two-fold increase in σ*_c_* with GG content from 0.5% to 1% by weight was observed with LHS-1 (□); similar trends were also observed for σ*_t_*, see solid lines in Fig. 4(b). In contrast, MGS-1 (△) showed a saturation in both σ*_c_* and σ*_t_* with GG content around 1%, see dashed lines in Fig. 4(b). The optimal GG content was hence determined to be around 2% and 0.5% for lunar and Martian regolith, respectively.

It is perhaps likely that this difference in σ*_c_*, σ*_t_* dependence on gum concentration is due to the nature of binding alluded to earlier. It is also important to note that for concentrations above 2%, material compaction during the PADC became significantly challenging. This precluded testing for concentrations above 2%. Water content in the pre-fill mixture is another important parameter— insufficient water can impede proper hydration and bonding, while excessive water can weaken the structure by increasing voids.

### 3.3 Failure mechanisms and crack propagation in BRCs

As discussed in Sec. 2, we used digital image correlation (DIC) to analyze failure in the tensile test samples. This is not done for the compressive samples because the sample thickness mandated by the test standard results in out-of-plane deformation. The results for three different gum concentrations (GG from 0.5% to 1.5%) are summarized in Fig. 5 and pertain to LHS-1 based BRC brick. The conclusions to be presented were found to be very similar for the other simulants as well.

**Figure 5:**
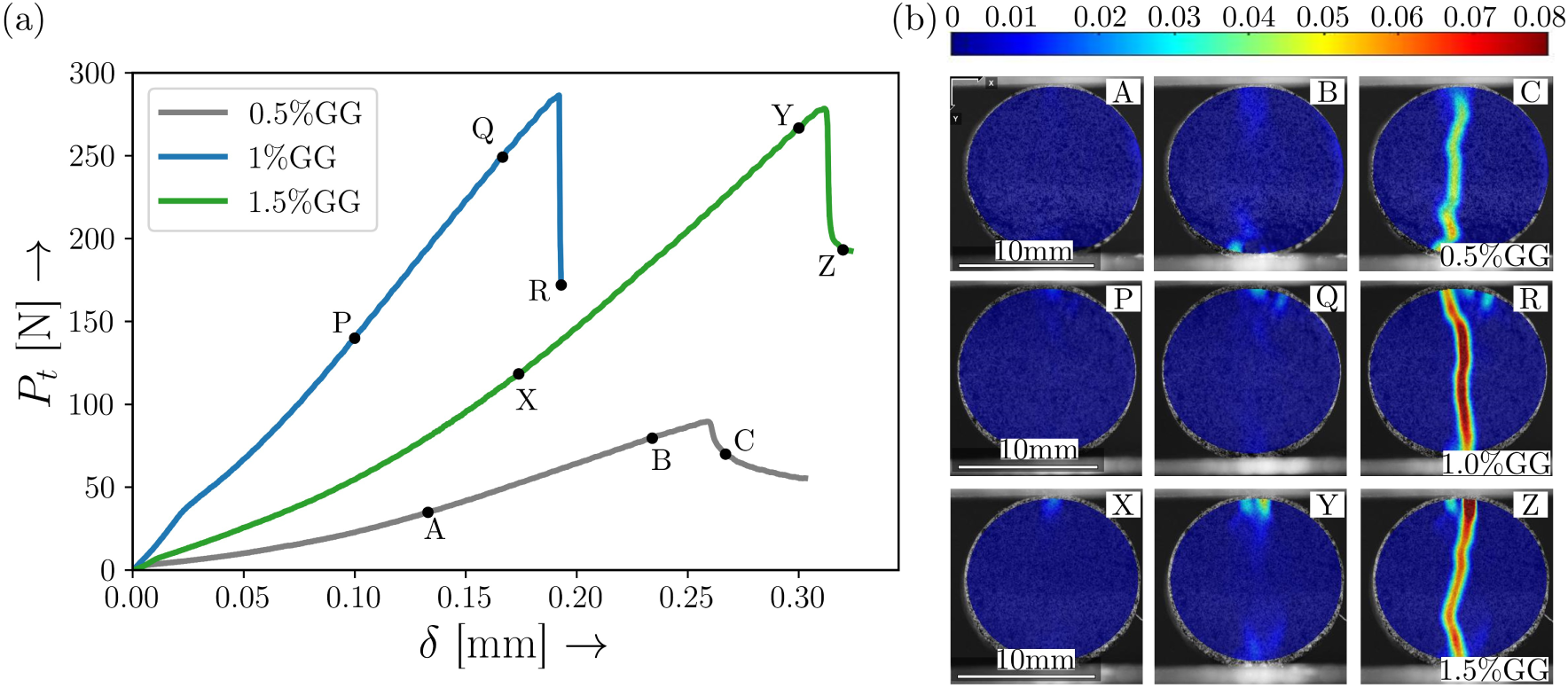
Failure of BRC bricks made from LHS-1 under tensile loading. (a) Load displacement graphs for three different guar gum concentrations, each with three points, corresponding to snapshots well before maximum load (A, P, X), close to maximum load (B, Y, Q), and just following failure (C, R, Z). The corresponding snapshots, with superimposed tensile strain fields (from DIC), are reproduced in panel (b). Color represents the Lagrangian strain transverse to the loading direction. Note that the failure pattern is unchanged by the binder concentration, and only the strain at failure is altered.

In panel (a) of this figure, three different loads (P*_t_*) vs. displacement (δ) graphs are presented, with varying GG concentrations as indicated in the legend. Specifically, each curve has three points marked—the first during deformation and well before failure (A, P, X), the second very close to the point of failure (B, Q, Y), and the third post-failure and load reduction (C, R, Z). Corresponding images of the brick, with tensile strain superimposed, are shown in panel (b). Here, the tensile strain component (ɛ*_xx_*, say) computed is along an axis (x) perpendicular to the loading direction (y). As a result, cracks correspond to large values of the ‘opening strain’ ɛ*_xx_*. Additionally, the strain fields are computed by comparing the current image with a reference image before deformation starts, and they correspond to the total (and not incremental) Lagrangian strain.

It is clear from the results for all three concentrations that failure inevitably occurs via the nucleation of a single crack at the loading platens (see images corresponding to points B, Q, Y) that then propagates largely linearly in the direction of loading, see images corresponding to points C, R, Z in panel (b). By comparing the data in panels (a) and (b) of Fig. 5, it is clear that the mode of failure is independent of the gum/ binder concentration and occurs via a single crack along the loading direction. However, the strain and, consequently, the load at which this occurs increases with concentration—the higher the binder concentration, the larger the strain and load at failure and, consequently, the larger the tensile strength of the sample.

### 3.4 What about other bio-polymer binders?

To compare the dependence of BRC strength on binder chemistry, we evaluated bricks produced via PADC with three different biopolymer binders—guar gum (GG), xanthan gum (XG), and acacia gum (AG)—at a fixed concentration of 1% w/w of the simulant (0.4 g gum powder in 40 g of simulant). Results of compressive σ*_c_* and tensile σ*_t_* strength evaluation of these samples are shown in Fig. 6. The error bars in this plot represent the standard deviation from four sample measurements.

**Figure 6:**
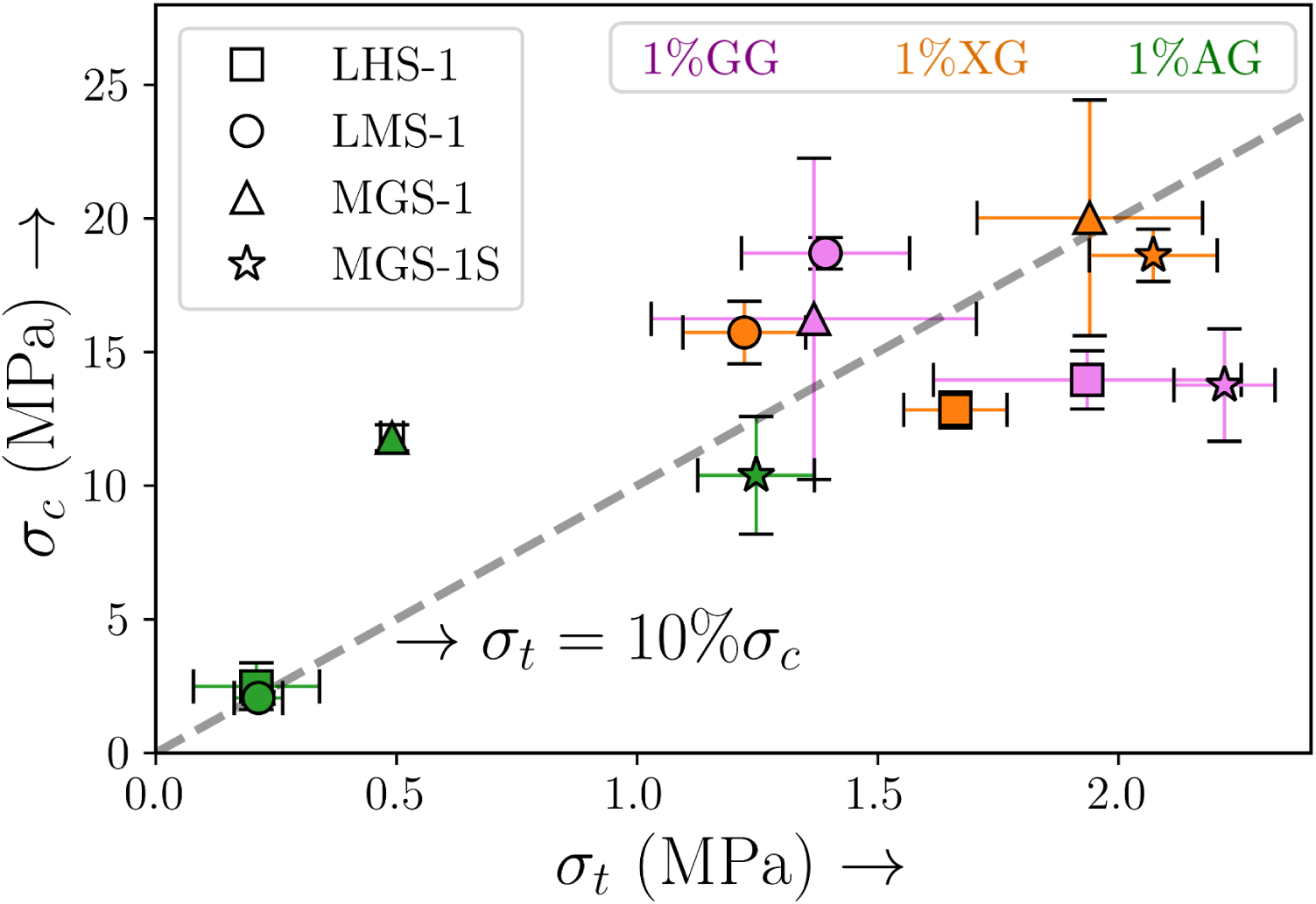
Scatter plot showing compressive (σ*_c_*) and tensile (σ*_t_*) strengths for different simulants (□ – LHS-1, ◦ – LMS-1, △ – MGS-1, and ⋆ – MGS-1S) mixed with guar gum (violet), xanthan gum (orange) and acacia gum (green), respectively. The marker shows the average strength, and the error bar represents the standard deviation. All samples were made using PADC (5 tons compaction), followed by 60*^◦^*C oven drying for 12 hours. The dashed black line represents the relationship σ*_t_* = 0.1σ*_c_*.

Upon comparing the strengths of the four different regolith simulants, it is clear that both GG and XG perform comparably with all simulants, and much better than AG. This is notable since GG and XG have been identified as being excellent binders owing to their high molecular weights, hydrophilicity, and stability under varying conditions [33, 40]. When mixed with water, XG forms a stable hydrophilic viscous colloid, while GG demonstrates thickening and gelling properties upon dissolution. Although AG is soluble in water, its binding and thickening capabilities are generally inferior to those of GG and XG. This enhancement is attributed to the coating and binding actions of polysaccharides on sand particles, which contribute to improved moisture retention within the regolith.

Finally, the dashed line in Fig. 6 represents σ*_t_* = 0.1σ*_c_*; it is seen that the strength values for all three polysaccharide gums appear to fall close to this line. This relationship is consistent with prior studies on terrestrial soils [47]; for instance, in concrete the tensile strength can range from approximately 7% to 15% of the compressive strength. This characteristic is essential because while concrete performs well under compressive loads, it is relatively weak in tension, notably leading to failure under bending loads.

### 3.5 Integrity of brick under simulated environment

For BRCs to be eventually usable in extra terrestrial environments, they must be able to endure large temperature fluctuations as well as low pressures. This endurance can be evaluated in two different ways—first, the effect of such environmental fluctuations on the pre-fill mixture itself, which determines the possibility of the process being used under these conditions. The second is to evaluate the resilience of the finally produced BRC bricks themselves to these fluctuations. We comment on both of these modalities in this section.

#### 3.5.1 Role of temperature fluctuations

Temperatures on the lunar and Martian surfaces range from -223*^◦^*C to 123*^◦^*C, and from -140*^◦^*C and 20*^◦^*C, respectively. It is clear that producing BRC consolidate bricks is not possible for this entire range; for instance, guar gum by itself can disintegrate beyond 80*^◦^*C. Hence, we evaluate the effect of large and small temperatures on already consolidated BRC bricks. Results are reported for LHS-1 with 1% GG using the PADC process; analogous results were obtained with the other simulants as well.

We subjected LHS-1 based bricks to a temperature of 175*^◦^*C in a hot air oven for 12 hours, as well as -80*^◦^*C in a deep freezer for 12 hours. This time duration was chosen to be concomitatnt with the time required for brick formation. Additional intermediate temperatures were also chosen—we selected soaking temperatures of -80*^◦^*C, -20*^◦^*C, 60*^◦^*C, 120*^◦^*C, and 175*^◦^*C. Following this soaking, the samples were subjected to mechanical testing to evaluate any changes in σ*_c_*, σ*_t_*, the results are summarized in Fig. 7. It is clear from this figure that the bricks continued to retain their room-temperature strength even after being subjected to these large temperature changes; both σ*_c_* and σ*_t_* did not change by more than a few percent, well within experimental error. It is therefore safe to conclude that, while the bricks have to be cast at room temperature to prevent degradation of the biopolymer constituents, once formed, they are capable of withstanding large temperature fluctuations with no measurable change in their strength. It is likely that the strong bonds and low porosity resulting from the PADC process are responsible for this robust strength.

**Figure 7:**
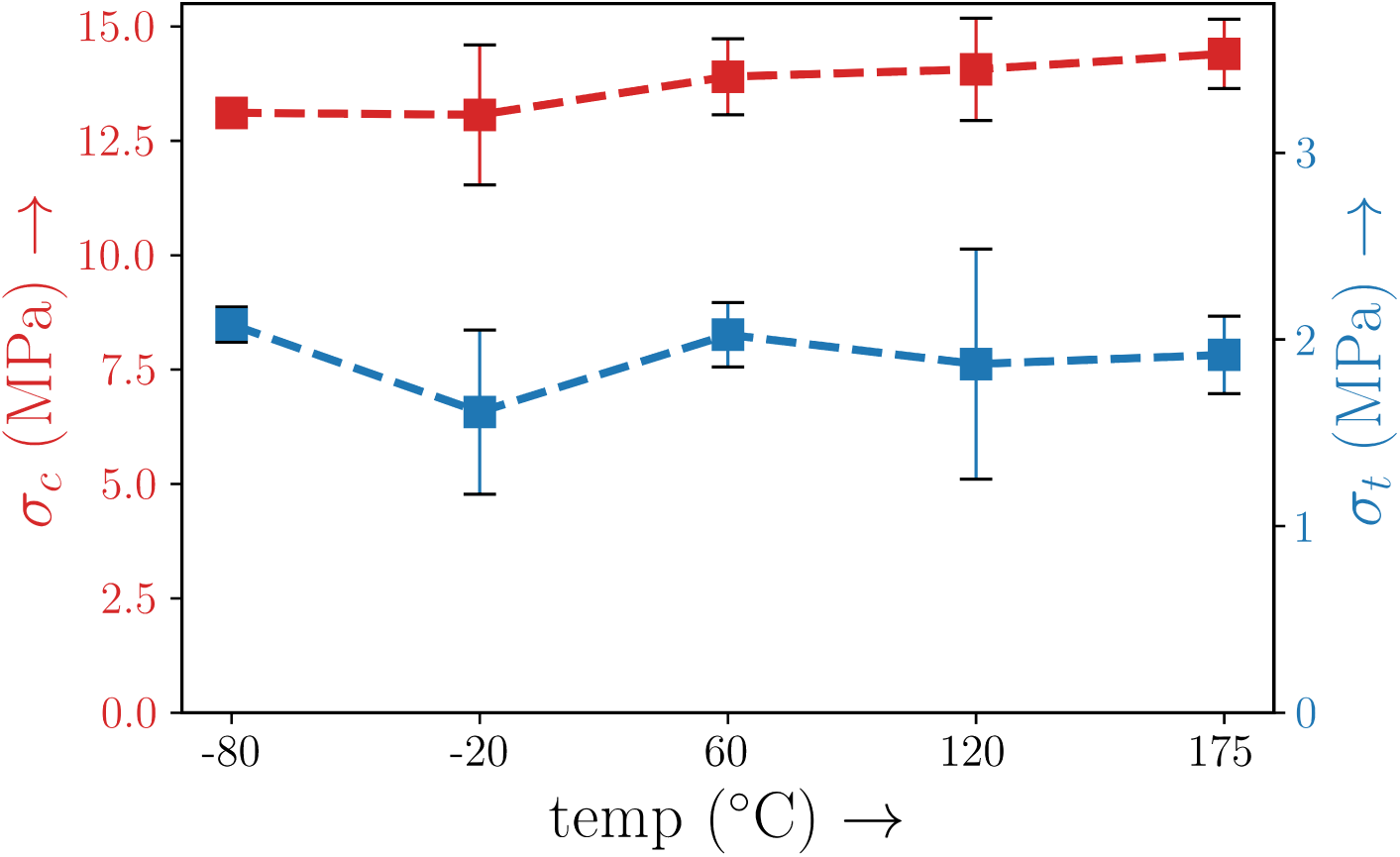
Variation in compressive σ*_c_* and tensile σ*_t_* strengths of LHS-1 based BRC bricks (1% w/w guar gum, PADC process) with temperature. Brick strength, post-casting, is seen to be largely unchanged by large temperature variations.

#### 3.5.2 Can we produce BRC in Vaccuum?

To now evaluate if the bricks can themselves be produced under vaccuum environments, we subjected SC consolidated bricks, prior to hardening, to a vaccuum environment, see Fig. 8(a). Given the varied chemistry of the simulants, including their water retention ability and presence of inherent binders, separate samples were prepared using LHS-1, LMS-1, and MGS-1 simulants, each combined with 0% and 1% guar gum (GG) and DI water, see Figs. 8(b) and (c), respectively. The pressure for consolidation was set to be ∼ 20 mbar (same as the pressure on the surface of Mars), in contrast with the consolidation performed at room temperature (1 bar), as discussed in Section 3.1. At this low pressure, the boiling point of water is significantly reduced to ∼ 15*^◦^*C, leading to water evaporation from within the pre-fill mixture. As a result, the samples completely collapsed in the absence of any binder, see corresponding images in panel (b). It is clear that the Martian simulant, with its inherent binder, was somewhat immune to this effect.

**Figure 8:**
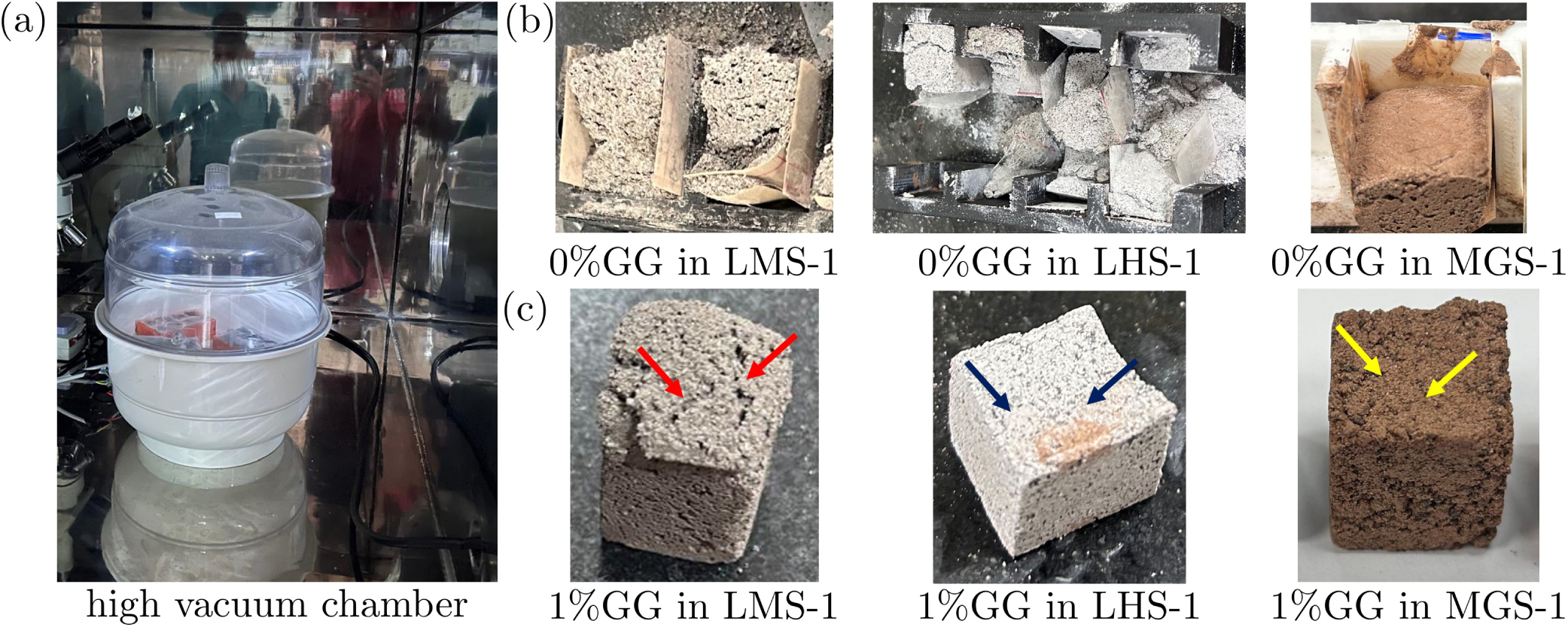
Effect of vaccuum conditions on BRC brick production and use. (a) Simple cast (SC) samples from LHS-1, LMS-1, and MGS-1 are placed inside a desiccator and then placed within a vaccuum chamber. Panels (b) and (c) show the SC specimens under low pressure (∼ 20 mbar) for LHS-1, LMS-1, and MGS-1 and with 0%, and 1% binder concentration, respectively.

On the other hand, the presence of GG binder appeared to minimize this effect, resulting in BRC consolidates that didn’t simply disintegrate, see panel (c). However, additional pores did form where water escaped the surface (see at arrows in each image). It is interesting that these marks were particularly pronounced with LMS-1 when compared with both MGS-1 and LHS-1. It thus appears that bricks can be produced even via the SC process under vaccuum conditions. The possibility of using PADC for minimizing water loss under vaccuum conditions is one that we could not evaluate in the present study. However, based on the earlier results, we believe that this should enable brick production under these extreme environments.

To evaluate performance of already formed BRC bricks (1% GG, PADC) under vaccuum conditions, all three bricks—LHS-1, LMS-1 and MGS-1—were subjected to a pressure ∼ 20 mbar for 24 hours. Visual inspection of the bricks before and after this exposure revealed no observable changes in the surface porosity or structure, see corresponding images in Fig. 9 for LHS-1 in particular. Following this soaking, the compressive strength of the bricks was again evaluated and the results are reproduced in table 3, in comparison with room pressure testing (1 bar). It is clear from these results that the bricks do not undergo any observable strength degradation after being subjected to vaccuum conditions. The use of 1% GG binders hence appears to result in bricks that can withstand both temperature and pressure fluctuations as long as they are produced within a controlled environment.

**Figure 9:**
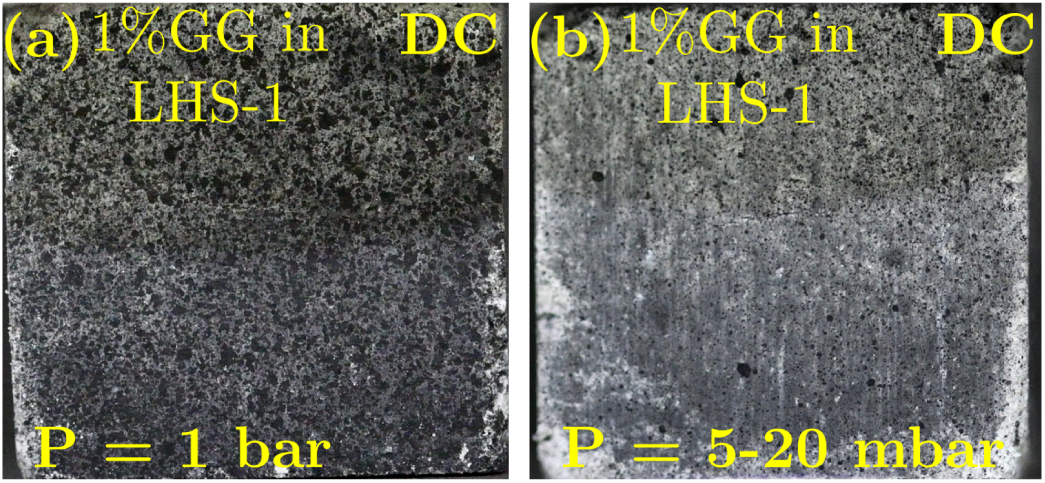
Comparison of the surface texture of LHS-1 PADC specimen, using the PADC process. (a) Surface prior to vaccuum exposure and (b) after being placed in a vaccuum for 24 hours.

**Table 3:** The compressive strength (σ*_c_*) of LHS-1, LMS-1, and MGS-1 consolidates mixed with 1% GG, exposed to vaccuum conditions (soaking pressure ranging from 5 to 20 mbar), compared to bricks tested under ambient pressure (1bar).

| Simulant | Soaking pressure (mbar) | $\sigma_c (MPa)$ |
| --- | --- | --- |
| LHS-1 | 1000 | $10.7 \pm 0.7$ |
| | 5-20 | $10.1 \pm 0.8$ |
| LMS-1 | 1000 | $19.9 \pm 1.8$ |
| | 5-20 | $16.7 \pm 0.4$ |
| MGS-1 | 1000 | $17.9 \pm 1.2$ |
| | 5-20 | $15.4 \pm 5.1$ |

## 4 Discussion and summary

Our study shows that polysaccharide biopolymers can be effectively used for binding regolith simulant into consolidated load-bearing structures. Specifically, the use of pressure-assisted dice casting (PADC) route can result in bricks—termed biopolymer regolith composite (BRC) bricks— with compressive strengths in excess of 15 MPa from both lunar and Martian simulants. This process is inherently more scalable, and, therefore, in line with the *in situ* resource utilization (ISRU) paradigm. It has minimal energy requirements, when compared to processes such as furnace (or kiln) sintering and is therefore much more easily deployed in remote and/or extraterrestrial environments.

The performance of lunar and martian simulants with the PADC process utilizing biopolymer binders is somewhat different due to their inherently varied chemistry. In the case of Martian simulants, optimal strength is reached for guar gum concentrations of around 0.5%, whereas in lunar simulants, this value is close to 2%. The application of pressure in the PADC process precludes the use of higher binder concentrations with both types of simulants. It is expected that the presence of inherent binders in Martian simulants results in slightly larger compressive and tensile strengths compared to lunar simulants. Furthermore, a change in binder concentration does not fundamentally change the failure mechanisms in BRC bricks, but only the strain (or, consequently, stress) at failure. This was confirmed via simultaneous force measurements and DIC analysis.

Our results have also shown that other biopolymers from the polysaccharide family, such as xanthangum can also efficiently bind regolith simulants via the PADC process. While the process requires some temperature and pressure control (to prevent binder degradation), the final bricks themselves can be subjected to significant temperature and pressure variations, with little to no change in their compressive or tensile strengths. We believe that this makes BRC-based bricks a suitable candidate for use in harsh extraterrestrial environments while adhering to the guidelines of the ISRU paradigm.

## Acknowledgments

The authors acknowledge financial support (AK for DST/TDT/AM/2022/345(G)), and (KV for DST/TDT/AM/2022/280(G)) from the Department of Science and Technology–DST, Govt. of India, under the Advanced Manufacturing Technologies program, for carrying out this work.

